# Cerebellar degeneration impairs strategy discovery but not strategy recall

**DOI:** 10.1101/2022.10.07.511309

**Authors:** Jonathan S. Tsay, Lauren Schuck, Richard B. Ivry

**Author notes:** **Corresponding author Information:** Name: Jonathan Tsay & Lauren Schuck, &, Address: 2121 Berkeley Way, Berkeley, CA 94704. These authors contributed equally to this work (Jonathan S. Tsay, Lauren Schuck).

## Abstract

The cerebellum is recognized to play a critical role in the automatic and implicit process by which movement errors are used to keep the sensorimotor system precisely calibrated. However, its role in other learning processes frequently engaged during sensorimotor adaptation tasks remains unclear. In the present study, we tested the performance of individuals with cerebellar degeneration on a variant of a visuomotor adaptation task in which learning requires the use of strategic re-aiming, a process that can nullify movement errors in a rapid and volitional manner. Our design allowed us to assess two components of this learning process, the discovery of an appropriate strategy and the recall of a learned strategy. Participants were exposed to a 60° visuomotor rotation twice, with the initial exposure block assessing strategy discovery and the re-exposure block assessing strategy recall. Compared to age-matched controls, individuals with cerebellar degeneration were slower to derive an appropriate aiming strategy in the initial Discovery block but exhibited similar recall of the aiming strategy during the Recall block. This dissociation underscores the multi-faceted contributions of the cerebellum to sensorimotor learning, highlighting one way in which this subcortical structure facilitates volitional action selection.

## Introduction

Motor adaptation ensures that well-learned movements remain accurate across a broad range of contexts (1,2). For example, a boxer fights against fatigue to keep his arms up and maintains a proper defensive posture, while a sailor adjusts her sails in response to variations in the wind.

Adaptation does not rely on a unitary process, but instead engages multiple learning mechanisms. A basic dichotomy delineates these processes into two general categories (1–9), those that entail volitional strategic changes associated with action selection (10) and those that are implicit, operating in an automatic way to ensure the sensorimotor system is properly calibrated (11–14). To illustrate this dichotomy, Stephen Curry can volitionally adjust his shooting angle to avoid an opponent seeking to block his shot, while subconsciously and automatically fine-tune the effort required to maintain a stable shooting angle in response to muscular fatigue.

It is clear from previous studies that the cerebellum plays a key role in implicitly recalibrating the sensorimotor system in response to force field and visuomotor perturbations (9,15–20). Implicit recalibration is severely attenuated in patients with cerebellar degeneration compared to their age-matched controls (11,15,17,20–23), and cerebellar lesions in non-human animals produce impairments in a wide range of adaptation tasks (24–26). Similarly, physiological correlates of implicit recalibration are prominent in the cerebellum (27–33).

More recently, researchers have begun to examine the contribution of the cerebellum to strategic processes engaged in motor adaptation. This work has generally involved variants of a visuomotor adaptation task in which participants move in a virtual reality setup where vision of the arm is occluded, and feedback is limited to a cursor. Adaptation processes are engaged by manipulating the relationship between the arm and cursor, usually by imposing a rotational transformation. While the resultant error will automatically engage implicit recalibration, participants are also likely to strategically adjust their aim to offset the perturbation, especially if the perturbation is large (e.g., aim in clockwise direction in response to a 60° counterclockwise rotation). Individuals with cerebellar degeneration (CD) are able to employ the appropriate re-aiming strategy if provided with verbal instructions and/or visual cues (34,35). In contrast, CD participants exhibit a marked deficit in re-aiming when required to discover the appropriate strategy (36).

Inspired by this dissociation across studies, we hypothesized that the cerebellum may be selectively involved in the *discovery* of a re-aiming strategy, rather than the *recall* of a known strategy. To test this, we recruited participants with CD and age-matched controls to participate in a visuomotor adaptation task, one designed to isolate these components of strategic re-aiming. Critically, the participants were exposed to the perturbation twice, with the initial exposure block assessing strategy discovery and the re-exposure block assessing strategy recall. A deficit in strategy discovery in the CD group will manifest as less effective re-aiming during the initial exposure block compared to their age-matched controls, whereas a deficit in strategy recall will manifest as less effective re-aiming during the re-exposure block. Taken together, these two probes will specify *how* the cerebellum may contribute to strategic use during sensorimotor learning.

## Materials and Methods

### Ethics Statement

All participants gave written informed consent in accordance with policies approved by the UC Berkeley Institutional Review Board. Participation in the study was in exchange for monetary compensation.

### Participants

We recruited 16 individuals with a medical diagnosis of cerebellar degeneration (CD) from our clinical database. The database is composed of individuals from around the country who responded to online advertisements either posted on the National Ataxia Foundation website or distributed by leaders of local support groups. For those who agreed to participate, we conducted an online video interview to obtain a medical history, a neurological assessment of CD symptoms using the Scale for Assessment and Rating of Ataxia (SARA; (37)), and an evaluation of general cognitive status using the Montreal Cognitive Assessment (MoCA; (38,39)). Table 1 provides a summary of group demographics and Table S1 provides this information for each participant.

**Table 1:** CD and matched control participants. SARA scores can range from 0 to 40 (where higher scores indicate greater symptomology). MoCA scores can range from 0 to 30 (where lower scores indicate greater impairment). Mean ± SD is provided.

| Group | N | Age | Sex | Handedness | Years of Education | MoCA | SARA |
| --- | --- | --- | --- | --- | --- | --- | --- |
| CD | 16 | 57.5 ± 11.8 | 5 M, 11 F | 15R, 1L | 16.9 ± 2.7 | 26.1 ± 2.1 | 14.3 ± 5.7 |
| Control | 16 | 59.8 ± 11.7 | 7 M, 9 F | 15R, 1L | 17.5 ± 2.0 | 27.7 ± 1.7 | – |

The sample size was based on an analysis of the data from a published study comparing strategic re-aiming in CD and age-matched controls (36). In that study, groups of 10 CD and control participants yielded a large effect size of 3.75 and power of .99. We opted to recruit 16 individuals for our groups as a conservative and feasible sample size.

Inclusion in the CD group was based on genetic confirmation of spinocerebellar atrophy (SCA) or a clinical diagnosis of ataxia. 9 of the 16 individuals had an identified subtype (SCA1: 1; SCA2: 2; SCA3: 5; SCA6: 1). One individual in the CD group was classified as SAOA (sporadic adult-onset ataxia). Six individuals did not have genetic testing (i.e., unknown etiology) but had MRI confirmation of cerebellar degeneration. Based on the medical history component of the online interview, we excluded candidate participants who self-reported ataxia but neither had MRI confirmation of cerebellar degeneration or genetic confirmation of SCA.

The SARA and MoCA were modified for online administration (40). For the SARA, two items were scored based on self-reports due to concerns about safety during remote evaluation (Gait and Stance). For the MoCA, we eliminated the Alternating Trail Making Task in the online version because it requires a paper copy. The scores were rescaled to the standard score range (0 – 30). As shown in Table 1, the CD group exhibited moderately severe cerebellar ataxia (a SARA score greater than 14) (37). Five of the CD participants exhibited mild cognitive impairment (a MoCA score less than 26) (38). We opted to include these participants given our general interest in the contribution of the cerebellum to cognition, and here in this study, with a specific focus in testing the impact of CD on strategic re-aiming.

We recruited a sample of 16 matched controls, all of whom do not have a history of neurological disorders. The CD and Control groups did not show significant differences in terms of Sex, Handedness, Age (*t*(30) = 0.5, *p* = 0.60, *D* = 0.2), Years of Education (*t*(30) = 0.8, *p* = 0.50, *D* = 0.3), and MoCA scores (t(30) = 1.6, *p* = 0.10, *D* = 0.6).

### Apparatus and General Procedures

Participants used their own laptop or desktop computer to access a customized webpage that hosted the experiment (41). To provide instructions and ensure task compliance, an experimenter was present through a video link during the one-hour experimental session. Participants used their computer trackpad or mouse to perform the reaching task (sampling rate typically ~60 Hz). The size and position of stimuli were scaled based on each participant’s screen size. For ease of exposition, the stimuli parameters reported below are for a typical monitor size of 13’’ with screen resolution of 1366 × 768 pixels (42).

Reaching movements were performed by using the computer trackpad or mouse to move the cursor across the monitor. Each trial involved a planar movement from the center of the workspace to a visual target. The center position was indicated by a white circle and the target location was indicated by a blue circle (both 0.5 cm in diameter). On the typical monitor, the radial distance from the start location to the target was 6 cm. The target appeared at one of two locations on an invisible virtual circle (60° = upper right quadrant; 210° = lower left quadrant). The movement involved some combination of joint rotations about the arm, wrist, and/or finger depending on whether the trackpad or mouse was used. In our prior validation work using this online interface and procedure, the exact movement and the exact device used did not impact measures of performance or learning on visuomotor adaptation tasks (41).

To initiate each trial, the participant moved their cursor to the circle positioned at the center of the screen, the start location. The cursor was only visible when the white dot was within 2 cm of the start location. This minimized adaptation that might occur in response to feedback from the cursor during the return movement to the start location. Once the participant maintained their position in the starting circle for 500 ms, a blue circle would appear at one of the two target locations. The participant was instructed to reach after they heard the go-cue (i.e., an auditory “beep”), attempting to make the cursor land in the target. To minimize demands on amplitude control and/or movement termination, they were instructed to reach past the target.

We imposed a delay, randomly selected from rectangular distribution ranging from 1100-1300 ms between the target appearance and the auditory go-cue. By imposing a delay between target presentation and movement initiation, we sought to nullify any group differences in the time required for movement preparation and, as such, eliminate potential speed-accuracy differences. Participants heard the message “Wait for the tone!” if they initiated a movement prior to the go-cue and heard the message “Move earlier!” if they initiated a movement 800 ms after the tone.

The visual feedback cursor during the center-out movement could take one of three forms: Congruent feedback, rotated feedback, and no-feedback. For the congruent and rotated conditions, the cursor appeared at a single location (“endpoint feedback”). During congruent feedback trials, the angular position of the visual feedback was aligned with the direction of the hand movement. Thus, if the participant’s movement sliced through the target, the feedback appeared at the target location. During rotated feedback trials, the angular position of the cursor was rotated 60° relative to the position of the participant’s hand. Thus, if the participant’s movement sliced through the target, the position of the feedback cursor was shifted 60° along an invisible circle with a circumference equal to the target amplitude. The direction of this offset was either clockwise or counterclockwise (counterbalanced across participants and targets). During no-feedback trials, the cursor was extinguished as soon as the hand left the start circle and remained off for the entire movement.

Critically, when visual feedback was provided (congruent or rotated), we imposed an 800 ms delay between the time at which the amplitude of the movement reached the target distance and onset of the visual feedback. By delaying the visual feedback, we sought to eliminate the contribution of implicit processes that occur during motor adaptation, and as such, isolate the contribution of strategic re-aiming (43–46). The feedback cursor remained visible for 500 ms.

### Experimental Design

The experiment consisted of three phases. First, participants watched a video that introduced the various manipulations in the task, emphasizing that the primary goal was to hit the target with their visual cursor. The video also highlighted the use of a delayed go-cue and delayed feedback. Second, to become familiar with the web-based reaching environment as well as the congruent and no-feedback conditions, the participants completed 15 practice trials. Third, participants completed the main task, which entailed six blocks (80 movement cycles x 2 targets = 160 trials total): A baseline veridical feedback block (5 cycles), rotated contingent feedback block (i.e., “Discovery” block; 30 cycles), an initial no-feedback aftereffect block (5 cycles), veridical feedback block (5 cycles), rotated contingent feedback block (i.e., “Recall” block; 30 cycles), and a second no-feedback aftereffect block (5 cycles).

Before the start of each veridical feedback block, participants were provided the following instructions: “Please move your white cursor directly to the blue target immediately after the tone.” Before the start of each rotated contingent feedback block, participants were provided the following instructions: “Your white cursor will be offset from where you move. Hit the blue target with your white cursor.” Before the start of each no-feedback aftereffect block, participants were provided the following instructions: “Your white cursor will be hidden and no longer offset from where you moved. Please move directly and immediately to the blue target after the tone.”

### Data Analysis

All data and statistical analyses were performed in R. The primary dependent variable was the endpoint hand angle on each trial, defined as the angle of the hand relative to when the movement amplitude reached a 6 cm radial distance from the start position. To aid visualization, the data were collapsed across the two movements in a cycle (one per target). Data points were considered outliers when the hand angle deviated from a 5-trial trendline by more than 3 standard deviations. This resulted in the exclusion of 3.2 ± 1.9% (Mean ± SD) of the data points from the Control group and 2.7 ± 1.6% of the data points in the CD group.

We compared hand angle between groups in three *a priori* defined epochs (47): Early adaptation, late adaptation, and aftereffect. Early adaptation was defined as the initial ten movement cycles after the rotation was introduced (Discovery block: cycles 7 – 16; Recall block: cycles 47 – 56). Late adaptation was defined as the final ten movement cycles of the rotation blocks (Discovery block: cycles 31 – 40; Recall block: cycles 66 – 75). Aftereffect was defined as all movement cycles without visual feedback after the rotation was removed (following the Discovery block: cycles 41 – 45; following the Recall block: cycles 76 – 80).

Reaction time was defined as the time between the go-cue and the start of movement, with the latter operationalized as the time at which the hand movement exceeded 1 cm. Movement time was defined as the time from the initial movement to the time when the cursor reached 6 cm. Reaction time (Control: 499.4 ms [182.4, 747.5]; CD: 469.8 ms [-53.3, 1043.6]; *t*(29) = 0.7, *p* = 0.5, *D* = 0.2) and movement time (Control: 499.4 ms [115.5, 1167.8]; CD: 410.5 ms [120.5, 1005.5]; *t*(29) = 0.7, *p* = 0.5, *D* = 0.3) did not significantly differ between groups.

We employed F-tests with the Satterthwaite method to evaluate whether the coefficients (i.e., beta values) obtained from the linear mixed effects model were statistically significant (R functions: lmer, lmerTest, ANOVA). Pairwise post-hoc two-tailed t-tests (or Wilcoxon sign-ranked test when parametric assumptions were violated) were employed to evaluate whether hand angle and kinematic measures differed between groups. P-values were adjusted for multiple comparisons using the Tukey method. The degrees of freedom were also adjusted when the variances between groups were not equal. Standard effect size measures are also provided (*D* for between-participant comparisons; *D* for within-participant comparisons; 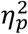 for between-subjects ANOVA) (48).

## Results

How does the cerebellum contribute to strategic re-aiming? To examine this, we tested participants with cerebellar degeneration (CD) and age-matched controls (N = 16/group) on a visuomotor rotation task designed to require the use of an aiming strategy. After an initial veridical feedback baseline block to familiarize participants with the task environment, the feedback cursor was rotated by 60° (Figure 1a). To compensate for this rotation, both groups exhibited significant hand angle changes in the opposite direction of the rotation, drawing the cursor closer to the target. When participants were asked to forgo their strategies and re-aim back to the target during the initial no-feedback aftereffect block, both groups were able to “switch-off” their strategies and successfully re-aim back to the target. Both groups exhibited minimal aftereffects, confirming that our delayed feedback manipulation was successful in eliminating implicit recalibration (43,44). Upon re-exposure to the rotation in the Recall block, participants were able to recall their aiming strategy, and subsequently switch back to aiming directly to the target in the second no-feedback aftereffect block. In sum, the adaptive changes observed in response to the 60° rotation were the result of strategic re-aiming, rather than implicit recalibration (also see Supplemental Materials).

**Figure 1:**
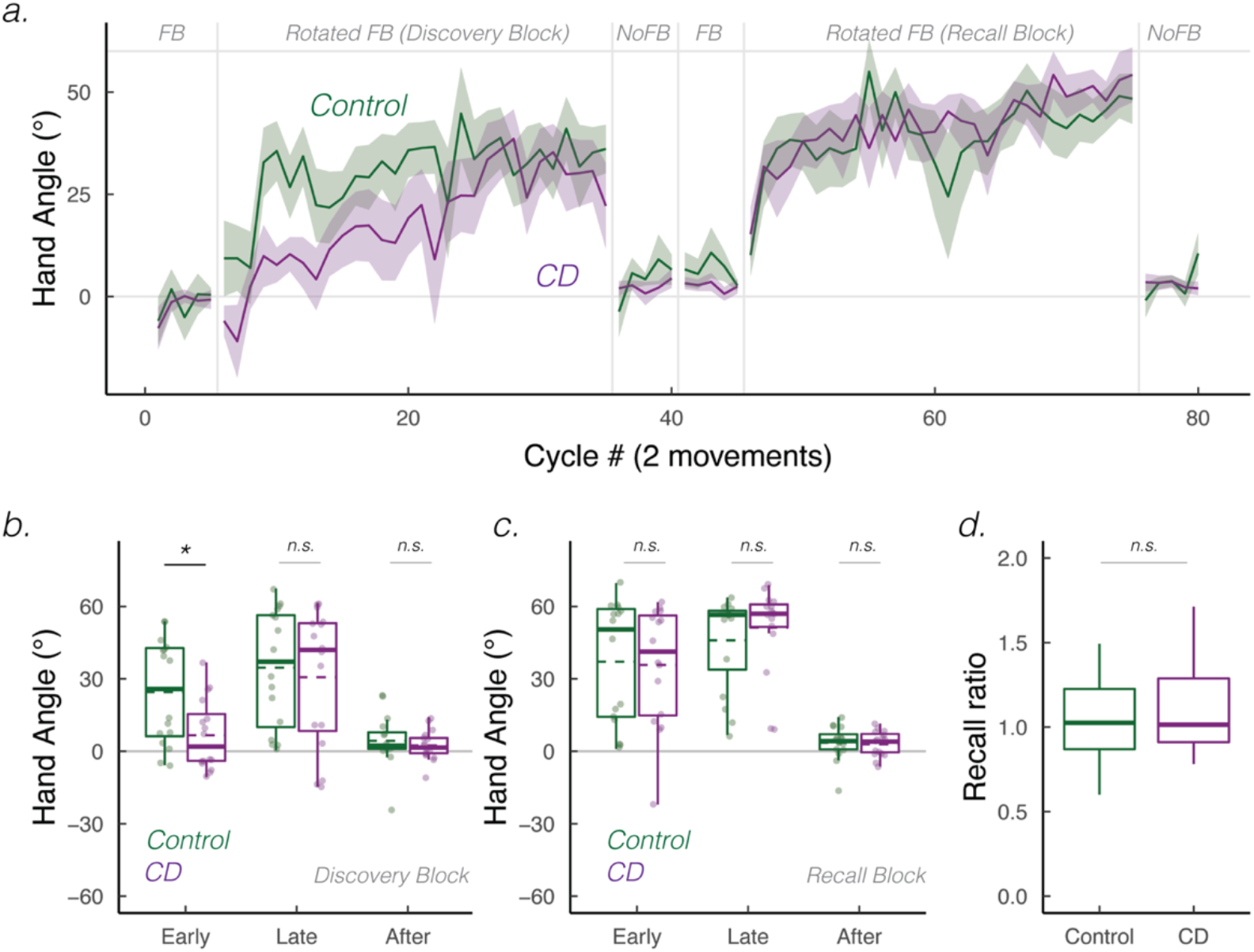
Cerebellar degeneration impairs strategy discovery but not strategy recall. **(a)** Mean time courses of hand angle for Control (dark green) and CD groups (dark magenta). Hand angle is presented relative to the target (0°). **(b)-(c)** Box plot denotes median hand angles (solid line), mean hand angles (dashed lines), 1^st^/3^rd^ IQR, and min/max during the three epochs of the **(b)** Discovery rotation block and **(c)** Recall rotation block. **(d)** Recall ratio, quantifying the degree to which late adaptation in the Discovery block was recalled during early adaptation of the Recall block. n.s. denotes that the group comparison between groups was not significant. * denotes p < 0.05. Translucent dots denote individual participants.

To statistically evaluate performance, we focused on three *a priori* defined epochs: Early, late, and aftereffect epochs (Figure 1b-c). For Control participants, hand angles deviated significantly from baseline to nullify the rotation during both early adaptation (24.5° [13.0°, 35.9°]; *t*(57) = 4.3, *p* < 0.001, *D*_*z*_ = 1.1) and late adaptation in the Discovery block (34.5° [23.0°, 45.9°]; *t*(57) = 1.2, *p* = 0.25, *D*_*z*_ = 1.3). For the CD participants, hand angles did not significantly deviate from baseline during early adaptation (6.6° [-4.8°, 18.1°]; *t*(57) = 1.2, *p* = 0.25, *D*_*z*_ = 0.4), evidence of a deficit in strategic discovery. The CD participants did exhibit significant adaptation during the late epoch of the Discovery block (27.8° [16.4°, 39.3°]; *t*(57) = 4.9, *p* < 0.001, *D*_*z*_ = 0.9). There was a main effect of Block for both early (*F*(1, 30) = 5.0, *p* = 0.03, 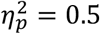) and late epochs (*F*(1, 30) = 4.3, *p* = 0.05, 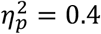), demonstrating that hand angles were greater in the Recall Block compared to Discovery Block.

Turning to our main question, we asked whether CD impacts strategy discovery and/or strategy recall. During early adaptation, there was a main effect of Group (*F*(1, 49) = 5.2, *p* = 0.03, 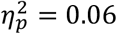) and a marginal interaction between Group and Block (*F*(1, 30) = 4.3, *p* = 0.05, 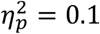). As can be seen in Figure 1b-c, the CD group showed attenuated early adaptation during the Discovery block compared to the Control group, but a similar level of adaptation during the Recall block. This inference was confirmed in a post-hoc comparison of the two groups for each of the adaptation blocks (Discovery block: (*t*(26) = 2.3, *p* = 0.02, *D* = 1.0; Recall block: (*t*(26) = 0.2, *p* = 1, *D* = 0.1). Interestingly, during late adaptation, we observed neither a main effect of Group (*F*(1, 44) = 0.63, *p* = 0.43, 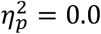) nor an interaction between these variables (*F*(1, 30) = 2.1, *p* = 0.16, 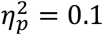). Thus, by the end of both rotation blocks, the CD and Control groups were equally successful in nullifying the visuomotor perturbation.

We recognize that focusing on hand angle may not be an appropriate measure of recall given that participants varied in terms of their strategy use over the course of the first rotation block. To account for this variation, we performed a second assay by calculating a “recall ratio”, quantifying the extent to which performance during the early adaptation epoch in the Recall block approached that observed in the late adaptation epoch of the Discovery block (i.e., the early recall epoch normalized by late discovery epoch). A recall ratio of 1 signifies complete strategic recall, below 1 signifies partial strategic recall, and above 1 signifies the use of a larger aiming strategy upon re-exposure to the visuomotor rotation. Based on this measure, both groups exhibited strong recall (Figure 1d): The Control group had a recall ratio of 1.3 (IQR: [0.9, 7.1]; Wilcoxon test against 0: *V* = 122, *p* = 0.003), and the CD group had a recall ratio of 1.0 (IQR: [0.8, 2.4]; *V* = 110, *p* = 0.03), respectively. Critically, the recall ratio did not differ significantly between groups (*W* = 135, *p* = 0.80), providing further evidence that strategic recall was not impaired in CD.

As shown in Figure 1c-d, both groups exhibited significant individual differences in their use of an aiming strategy. These individual differences may, in part, be driven by a generic cognitive deficit in a subset of the participants (i.e., inattention to the feedback, failure to understand the task). Indeed, there were some individuals whose late adaptation mean hand angles were close to baseline in both blocks; see (43,46)). These “non-learners” would also have non-sensical recall ratios since there was no learning to recall.

To address this concern, we defined “learners” as those participants who had a hand angle significantly greater than zero in the late adaptation epoch of the Discovery block (Figure 2a, data from 11 Controls and 10 CD participants). We then repeated our primary analyses on these subgroups, reasoning that group differences here would be related to impairment in the process of discovery and/or recall rather than a generic cognitive impairment. Our key comparisons were replicated: Adaptation in the CD group was selectively impaired in the early epoch of the Discovery block (Figure 2b; *t*(16) = 2.5, *p* = 0.04, *D* = 1.1). In contrast, the groups did not differ in all other epochs (Figure 2b-c; Discovery, late: *t*(19) = 0.9, *p* = 0.8, *D* = 0.4; Early: *t*(19) = 0.9, *p* = 0.40, *D* = 0.4; Late: *t*(19) = 0.1, *p* = 1, *D* = 0.0). The recall ratio also did not differ between groups (Figure 2d; *W* = 51, *p* = 0.80), altogether re-affirming that CD selectively impairs strategy discovery, but not strategy recall.

**Figure 2:**
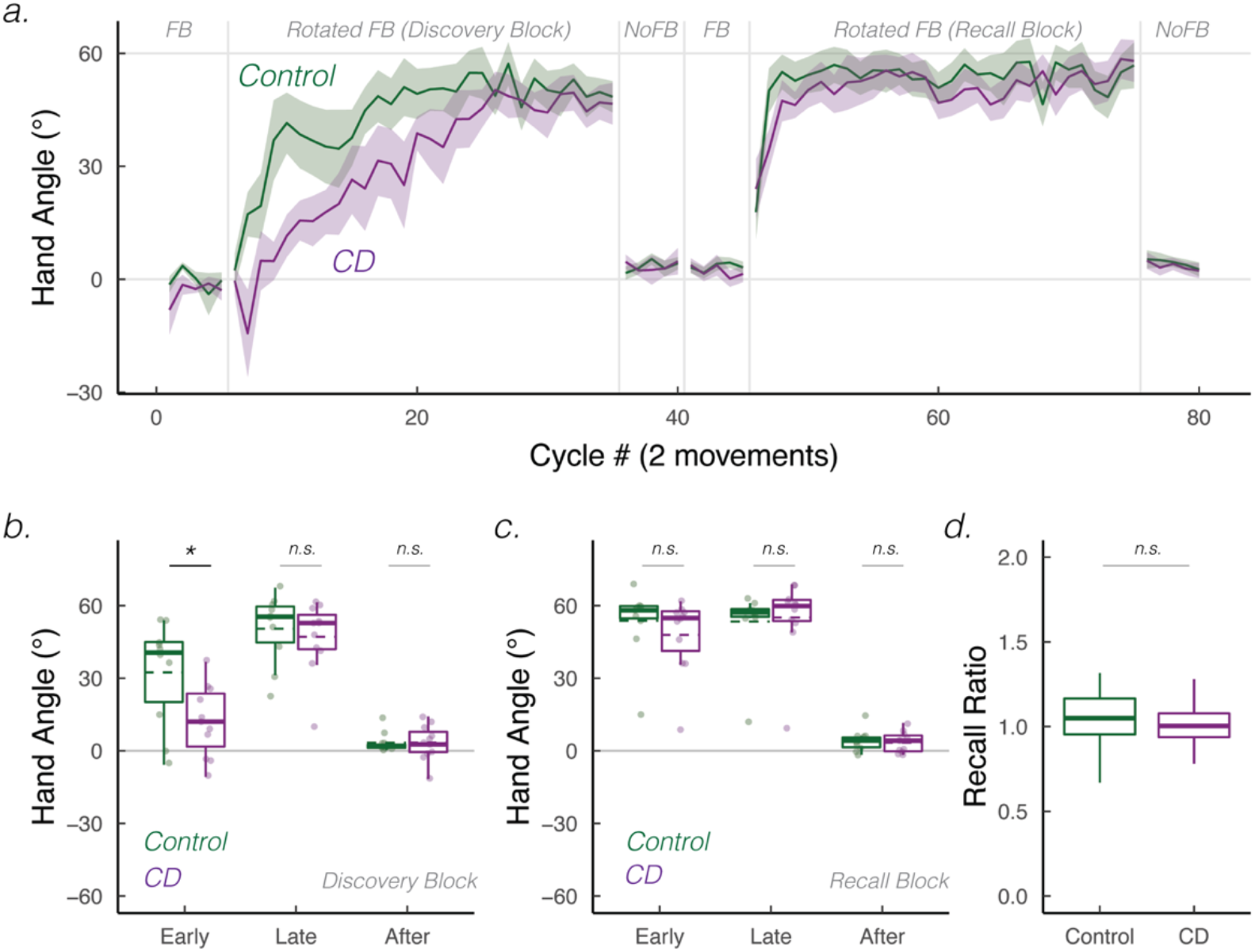
Verification of key findings when analyses were restricted to those showing evidence of strategy use in the Discovery block (“learners”). **(a)** Mean time courses of hand angle for Control (dark green) and CD groups (dark magenta). Hand angle is presented relative to the target (0°). **(b-c)** Box plot denotes median hand angles (solid line), mean hand angles (dashed lines), 1^st^/3^rd^ IQR, and min/max during the three epochs of the **(b)** Discovery rotation block and **(c)** Recall rotation block. **(d)** Recall ratio, quantifying the degree to which late adaptation in the Discovery block was recalled during early adaptation of the Recall block. n.s. denotes that the group comparison between groups was not significant. * denotes p < 0.05. Translucent dots denote individual participants.

## Discussion

While sensorimotor adaptation is often assumed to be automatic and implicit, motor adaptation tasks can also induce volitional and strategic re-aiming (48). Recent work has indicated that the integrity of the cerebellum is not only essential for implicit recalibration, but is also important for successful strategic re-aiming (50). Here we set out to ask *how* the cerebellum contributes to strategy use. We compared patients with cerebellar degeneration (CD) and matched controls on a visuomotor adaptation task in which performance changes were dependent on strategic re-aiming, using a design that allowed us to examine strategy discovery and strategy recall. Participants with CD exhibited attenuated early learning upon initial exposure to the visuomotor rotation, evidence of a strategy discovery impairment. However, the CD group eventually reached levels of learning similar to that of Controls. Moreover, the CD group were successful in recalling their strategy upon re-exposure to the rotation. Taken together, the results suggest that the cerebellum may play a selective role in strategy discovery but not strategy recall.

The cerebellum’s role in strategic re-aiming may parallel its role in cognition at large: One recent hypothesis centers on the cerebellum facilitating ‘mental simulations’ (47,51,52). For example, patients with CD exhibit impairments in covertly rotating a visual object held in mind or in covertly ‘moving’ along a mental number line during simple addition. Interestingly, strategy discovery in a visuomotor rotation task is also thought to require mental simulation, that is, the ability to mentally transform a motor plan that is initially oriented toward the target and then rotated to a direction aligned with the hypothesized aiming solution (53). A deficit in mental simulation may have impaired the CD group’s ability to rapidly discover a re-aiming strategy.

Could the discovery deficit in the CD group arise from impairments other than that of mental simulation? For example, the CD group may be impaired in estimating the direction and size of the rotation, reflecting a deficit in visual (53) and/or proprioceptive perception (i.e., perceiving the relative position of the visual cursor with respect to the moving hand) (54). That is, low-fidelity perceptual inputs ‘upstream’ may have impaired ‘downstream’ mental simulation. Future studies employing subjective reports of the perceived rotation size may unveil the stage in which the cerebellum contributes to strategy discovery (also see (55)).

What is clear is that degeneration of the cerebellum does not disrupt simple memory retrieval. For example, patients with CD were unimpaired on a task that required the retrieval of visual representations in working memory or in using a look-up table to perform simple multiplication (47). Correspondingly, previous visuomotor rotation studies have shown that CD participants do not show deficits in recalling an already discovered strategy or implementing a strategy provided through verbal instructions (55). The results of the present study are in accord with these findings in that the CD group was successful in recalling their re-aiming strategy upon re-exposure to the visuomotor rotation. Interestingly, their normal performance during recall (and at the end of the discovery block) would suggest that mental simulation is not required when a strategy is well-established. Successful recall in our current design could be achieved by memorizing two aiming locations, one for each target (56).

The inferences we can draw in terms of brain-behavior relationships is limited by the heterogeneity of our patient sample. Of note, our sample did include a large number of patients with SCA-3, a genetic variant that typically produces symptoms indicative of cerebellar and extra-cerebellar involvement (56) and is associated with depressed cognitive function (57). Note that 1) this subgroup did not stand out on our measures of strategy discovery or recall, suggesting that the impairment in strategy discovery is related to degeneration of the cerebellum; 2) the number of non-learners was similar in the CD and Control groups, arguing that the observed dissociation between discovery and recall is unlikely to be due to a general cognitive impairment in the CD group; and 3) our results have been corroborated by that of another unpublished dataset using a standard visuomotor rotation task (58). Nonetheless, revealing *which* cerebellar are critical for strategic discovery is an exciting area for future research, one that could be addressed in studies using larger samples to correlate changes in brain anatomy and re-aiming behavior or functional imaging studies to ask identify regions within the cerebellum associated with strategy discovery and how these regions interact with the rest of the brain, presumably to support this volitional process.

The current results add to the growing evidence that the functional contribution of the cerebellum in adaptation tasks extends beyond its well-established role in the implicit recalibration of the sensorimotor system. By using a task that minimizes the contribution of implicit recalibration, we provided clear evidence that CD disrupts strategic re-aiming. Importantly, the observed dissociation between strategy discovery and strategy recall points to how the cerebellum might contribute to strategic re-aiming, with the latter hypothesized to require the simulation of internal (50,59) and physical states (60,61). More generally, postulating constraints on the computational role of the cerebellum will be essential for advancing our understanding of how this subcortical structure interacts with the rest of the brain to support our motor and mental competences.

## Supplemental Materials

**Adaptive changes primarily arise from explicit re-aiming**: We sought to verify that the adaptive changes in the current experiment should be attributed to strategic re-aiming rather than implicit recalibration. To test this, we examined the hand angles during the two aftereffect blocks. After the Discovery block, aftereffects did not significantly deviate from baseline in both groups (Control: 4.3° [-1.5°, 10.1°]; *t*(15) = 1.6, *p* = 0.10, *D* = 0.4; CD: 2.4° [-1.0°, 5.8°]; *t*(15) = 1.5, *p* = 0.20, *D* = 0.4). After the Recall block, the Control group again exhibited no aftereffects (3.5° [-0.3°, 7.4°]; *t*(15) = 1.9, *p* = 0.07, *D* = 0.5). The CD group exhibited a small aftereffect (2.9° [0.1°, 5.7°]; *t*(15) = 2.2, *p* = 0.04, *D* = 0.6), a value that is negligible compared to typical aftereffects observed in the literature (15° - 25°) (11–13,62). Aftereffects neither differed significantly between Blocks (main effect of Block: *F*(1, 30) = 0.1, *p* = 0.75, 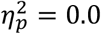) nor Groups (main effect of Group: *F*(1, 58) = 0.5, *p* = 0.48, 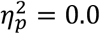; Group x Block interaction: *F*(1, 30) = 0.2, *p* = 0.71, 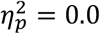). In summary, these results indicate that the delayed endpoint feedback manipulation successfully minimized the extent of implicit recalibration. By inference, we assume that the adaptive changes arise mainly from strategic re-aiming (63).

**Table S1:** Demographics for the individual participants in the group with cerebellar degeneration (CD) and the age-matched Control group. Participant Identification, Age, Sex, Handedness, scores for the Montreal Cognitive Assessment (MoCA), and Years of Education (YOE) are reported for all participants. MoCA scores ≥ 25 indicate no cognitive impairment. Scores for the Scale for Assessment and Rating of Ataxia (SARA) and the subtype of cerebellar degeneration are reported for the CD group. Higher SARA scores indicate more severe cerebellar symptoms and motor deficits.

| Cerebellar Degeneration (n = 16) |  |  |  |  |  |  |  | Control (n = 16) |  |  |  |  |  |
| --- | --- | --- | --- | --- | --- | --- | --- | --- | --- | --- | --- | --- | --- |
| ID | Age | Sex | Hand | Subtype | MoCA | YOE | SARA | ID | Age | Sex | Hand | MoCA | YOE |
| 1 | 59 | M | R | SCA3 | 28 | 16 | 5.5 | 1 | 71 | M | R | 24 | 18 |
| 2 | 51 | F | R | SCA2 | 25 | 16 | 12 | 2 | 49 | M | R | 29 | 20 |
| 3 | 50 | M | R | SCA3 | 26 | 19 | 11 | 3 | 60 | F | R | 24 | 16 |
| 4 | 50 | F | R | SCA1 | 29 | 20 | 10.5 | 4 | 38 | M | L | 30 | 18 |
| 5 | 65 | F | L | SCA2 | 26 | 16 | 14.5 | 5 | 62 | F | R | 28 | 20 |
| 6 | 64 | M | R | Sporadic SCA | 22 | 16 | 12 | 6 | 80 | M | R | 29 | 14 |
| 7 | 79 | F | R | Unknown | 28 | 18 | 11 | 7 | 61 | F | R | 29 | 17 |
| 8 | 79 | F | R | Unknown | 29 | 12 | 21.5 | 8 | 60 | F | R | 27 | 16 |
| 9 | 38 | F | R | Unknown | 26 | 12 | 23.5 | 9 | 58 | M | R | 29 | 15 |
| 10 | 57 | F | R | Unknown | 28 | 14 | 22.5 | 10 | 47 | M | R | 27 | 16 |
| 11 | 58 | F | R | Unknown | 25 | 17 | 18 | 11 | 69 | F | R | 28 | 18 |
| 12 | 44 | F | R | SCA3 | 29 | 18 | 4.5 | 12 | 44 | F | R | 28 | 18 |
| 13 | 54 | F | R | Unknown | 28 | 18 | 11 | 13 | 50 | F | R | 27 | 22 |
| 14 | 60 | M | R | SCA3 | 25 | 18 | 15.5 | 14 | 74 | F | R | 29 | 18 |
| 15 | 43 | F | R | SCA3 | 28 | 18 | 20 | 15 | 62 | M | R | 27 | 18 |
| 16 | 69 | M | R | SCA6 | 24 | 22 | 15.5 | 16 | 71 | F | R | 28 | 18 |

